# ZnJ6 is a DnaJ-like Chaperone with Oxidizing Activity in the Thylakoid Membrane in *Chlamydomonas reinhardtii*

**DOI:** 10.1101/2020.09.14.296681

**Authors:** Richa Amiya, Michal Shapira

## Abstract

Assembly of photosynthetic complexes is sensitive to changes in light intensities, drought, and pathogens that induce a redox imbalance, and require a variety of substrate-specific chaperones to overcome the stress. Proteins with cysteine (C) residues and disulfide bridges are more responsive to the redox changes. This study reports on a thylakoid membrane-associated DnaJ-like protein, ZnJ6 (ZnJ6.g251716.t1.2) in *Chlamydomonas reinhardtii*. The protein has four CXXCX(G)X(G) motifs that form a functional zinc-binding domain. Site-directed mutagenesis (Cys to Ser) in all the CXXCX(G)X(G) motifs eliminates its zinc-binding ability. In vitro chaperone assays using recombinant ZnJ6 confirm that it is a chaperone that possesses both holding and oxidative refolding activities. Although mutations (Cys to Ser) do not affect the holding activity of ZnJ6, they impair its ability to promote redox-controlled reactivation of reduced and denatured RNaseA, a common substrate protein. The presence of an intact zinc-binding domain is also required for protein stability at elevated temperatures, as suggested by a single spectrum melting curve. Pull-down assays with recombinant ZnJ6 revealed that it interacts with oxidoreductases, photosynthetic proteins (mainly PSI), and proteases. Our *in vivo* experiments with *Chlamydomonas reinhardtii* insertional mutants (ΔZnJ6) expressing a low level of ZnJ6, suggested that the mutant is more tolerant to oxidative stress. In contrast, the wild type has better protection at elevated temperature and DTT induced stress. We propose that DnaJ-like chaperone ZnJ6 assists in the prevention of protein aggregation, stress endurance, and maintenance of redox balance.

**One-sentence summary:** ZnJ6 is a redox-regulated DnaJ-like chaperone associated with the thylakoid membrane and involved in the prevention of protein aggregation and stress endurance.

## INTRODUCTION

Photosynthetic organisms are often challenged by biotic and abiotic stresses, resulting in a redox imbalance that must be counteracted by the organism to survive. The redox status of proteins in the chloroplast is mainly controlled and influenced by photosynthetic light reactions, which can lead to a rise in the generation of Reactive Oxygen Species (ROS) during stress. This occurs when the cells cannot dissipate excess of electrons due to an imbalance between the excited electrons and the carrier pathways (Erickson et al., 2015). Proteins containing cysteine residues and disulfide bridges are sensitive to redox changes that affect the redox status of these bridges, thus leading to structural changes, altered ability to function, and impaired ability to interact with partner proteins. Redox status of proteins, therefore, plays an essential role in cell signaling and anti-oxidizing defence (Klomsiri et al, 2011).

Here we focus on a novel thylakoid membrane-associated oxidase ZnJ6 (ZnJ6.g251716.t1.2) from *C. reinhardtii*. The protein contains four cysteine-rich CXXCX(G)X(G) motifs that form two C_4_ type zinc fingers. Given the similarity in this domain to that of DnaJ, ZnJ6 is categorized as a DnaJ-like protein (DnaJ E). It lacks all other motifs that are typical of DnaJ, such as the J and G/F domains, and there is also no homology to DnaJ in its C-terminus (Doron et al, 2018). To date, 20 proteins from this family were identified in *Arabidopsis* (Pulido and Leister, 2018), but their orthologs in *Chlamydomonas* were not yet determined, mainly due to their limited similarities. Many of the DnaJ-like proteins have a role as chaperones and in the assembly of photosynthetic complexes. One example is the Bundle Sheath Defective gene (BSD2) that was initially identified in maize as required for Rubisco biogenesis. (Brutnell 1990). BSD2 was extensively studied (Feiz et al., 2014) (Wostrikoff and Stern, 2007) and further verified as one of the five chaperones used for *in vitro* assembly of Rubisco (Aigner et al, 2017). Other examples are the thylakoid associated DnaJ-like PSA2 and LQY1 proteins, which interact with components of the PSI and PSII complexes, respectively (Fristedt et al, 2014)(Lu et al, 2011). Although a phylogenetic analysis identified ZnJ6 as the closest ortholog of the Maize BSD2 (Doron et al, 2018), here we show that ZnJ6 has a transmembrane domain and localizes in the thylakoid membrane, thus suggesting that it could have a different role.

ZnJ6 exhibits chaperone function, like other members of the DnaJ-like family of chloroplast proteins (Doron et al, 2018). We show that ZnJ6 has a protective role in preventing aggregation *in vitro* of Citrate Synthase (CS), a thermal sensitive protein target common in chaperone assays to examine protection against aggregation (Segal and Shapira, 2015). ZnJ6 protects CS regardless of its cysteine-rich domain. However, the cysteine dependent oxidative refolding ability of the protein was restricted to the recombinant wild type protein, and was not observed with its cys-mutant. The redox activity was established first by the Insulin aggregation assays, in which reduction opens the disulfide bonds that hold the two insulin subunits together, causing the β-subunit to precipitate. ZnJ6 was shown to prevent this precipitation. Another assay aimed to analyze the effect of ZnJ6 on the refolding of reduced denatured RNaseA (rdRNaseA), a redox-sensitive chaperone target. ZnJ6 assisted the native refolding and oxidation of rdRNaseA, thereby regaining its lost activity. However, the recombinant cys-mutant of ZnJ6 was not functional in both assays. Furthermore, the role of the zinc finger domain in providing protein stability at elevated temperature was established using a single spectrum melting curve.

To explore the potential role of ZnJ6, its interactome was examined as well. This analysis highlighted that ZnJ6 interacts with photosynthetic proteins, oxidoreductases, and proteases. To expand our understanding of the ZnJ6 function, we examined the role of the ZnJ6 *in vivo* using a *C. reinhardtii* insertional mutant (ΔZnJ6) that expresses a low level of the protein. When compared to wild type cells, ΔZnJ6 was more tolerant to oxidative stress caused by H_2_O_2_ and MeV, but appeared to be more sensitive to reducing conditions induced by DTT. The mutants also appeared to be sensitive to heat stress, with impaired growth and reduction in chlorophyll levels. Altogether ZnJ6 functions as a chaperone that also possesses oxidizing activity. It could therefore assist the cells in overcoming redox-related stress and possibly be involved in the assembly of the photosynthetic apparatus.

## RESULTS

### ZnJ6 from *C. reinhardtii* is localized in the Thylakoid membrane of the chloroplast

The localization of the protein in the cell can serve as the first indication for its potential function. For this, we determined the intracellular localization of ZnJ6 using biochemical sub-fractionation, followed by western analysis. Cytoplasmic fractions from a 1L culture were collected immediately after cell disruption by nitrogen cavitation and centrifugation, before chloroplast isolation. Chloroplasts were isolated using a Percoll step gradient. Isolated chloroplasts were washed and tested for the presence of intact chloroplasts. Thylakoid membranes were isolated from 250 ml of log-phase *Chlamydomonas* cells using a 3-step sucrose gradient (as described in the Materials and Methods section).

The isolated subcellular fractions, along with total cell protein, were subjected to western analysis using antibodies against marker proteins typical for each fraction. The cytoplasmic fraction was verified by its interaction with antibodies against HSP70A, and the chloroplast fraction was confirmed by its interaction with antibodies against the Oxygen evolving enzyme (OEE33) (Figure 1 A). ZnJ6 was in the chloroplast fraction. Next, membrane (M) and soluble (S) fractions of *C. reinhardtii* cells were also resolved over 12% SDS-PAGE. The membrane fraction was verified by antibodies against psbA, and the soluble fraction was verified by antibodies against HSP70A. Antibodies against Rubisco Activase showed that this protein was distributed between the membrane and soluble fractions. The presence of ZnJ6 in the membrane fraction was confirmed using specific antibodies raised against amino acids 1-165 (Figure 1B and supplemental figure S1). Finally, the thylakoid fraction was verified by antibodies against psbA. ZnJ6 was also shown to be in the thylakoid fraction (Figure 1C). This finding is supported by the presence of a predicted transmembrane domain in ZnJ6, as shown by TMHMM and Phobius servers (Supplemental Figure S2).

**Figure 1.**
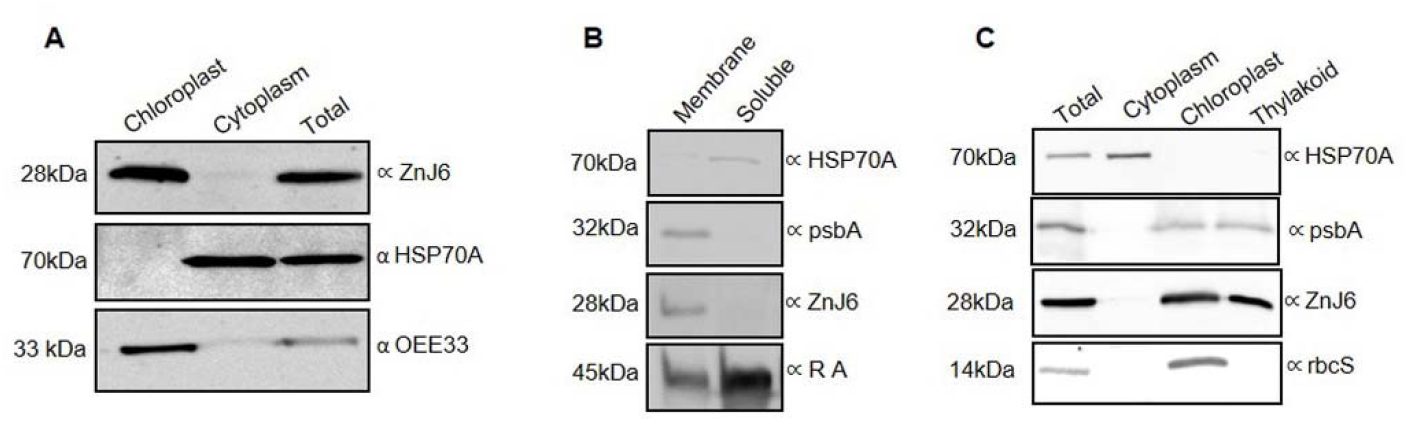
ZnJ6 is localized in the thylakoid membrane of the chloroplast. *C. reinhardtii* cells were grown until the late log phase and disrupted by nitrogen cavitation and further subfractionated. **A**, The purified chloroplasts, cytoplasm, and total proteins were subjected to western analysis using antibodies against OEE33 as a chloroplast marker and HSP70A as a cytoplasmic marker. **B**, The membrane and soluble fractions of *C. reinhardtii* cells were separated, and the presence of ZnJ6 in the membrane fraction was confirmed. psbA served as a membrane marker, and Rubisco Activase (RA) was is a marker that is distributed between the membrane and soluble fractions. **C**, Thylakoid membranes were isolated using a sucrose step gradient. Samples taken from the total extracts, cytoplasm, chloroplast, and thylakoid fractions were subjected to western analysis using antibodies against psbA as a thylakoid marker, rbcS as chloroplast markers, and HSP70A served as a cytoplasmic marker. Antibodies ZnJ6 specific antibodies unveiled its presence in the thylakoid membrane.

### CD analysis of affinity-purified ZnJ6 verifies that the recombinant protein is folded

To confirm whether the recombinant protein was folded, to examine its stability at high temperatures, and to test whether the Zn binding domain affected the folding, we measured the Circular Dichroism (CD) spectra of the recombinant wild type and cys-mutant proteins. Recombinant protein tagged with cleavable the Maltose Binding Protein tag (MBP) and the non-cleavable Streptavidin Binding Peptide (SBP). The protein was first purified over an amylose resin. The MBP tag was further cleaved from the fusion protein and the protein was further purified over Streptavidin resin (purified protein fraction was analysed over 12 % SDS gel, as shown in supplemental S3). 100 µl of the purified protein (with a concentration ≥ 100 µg/ml) was used for the analysis. Measurements were performed at a constant temperature of 23°C, at wavelengths ranging from 200-260nm (Figure 2A). In addition, measurements were taken at a constant wavelength of 222 nm with a temperature range from 20 °C to 80 °C, to examine whether elevated temperatures affected the protein structure. The results indicated that both the wild type and cys mutant proteins were folded (Figure 2A). However, we monitored differences in their stability at higher temperatures, depending on the presence or absence of the zinc-binding motif (Figure 2B). The melting curves show that both proteins remained folded at temperatures up to 65 °C (midpoint of transition state). However, the slope of transition between the folded and unfolded states was gradual with the ZnJ6 cys-mutant, unlike the wild type protein. This difference indicates reduced co-operative interactions in the mutant protein as compared to the wild type protein (Figure 2B). The stabilizing effect of the Zn binding domain was observed at higher temperatures, as the secondary structure of both proteins remained largely unaffected at optimum temperatures, regardless of the mutation in the zinc-binding motif. Thus, the cysteine-rich zinc motif was responsible for stabilization and folding of the structure at elevated temperatures.

**Figure 2.**
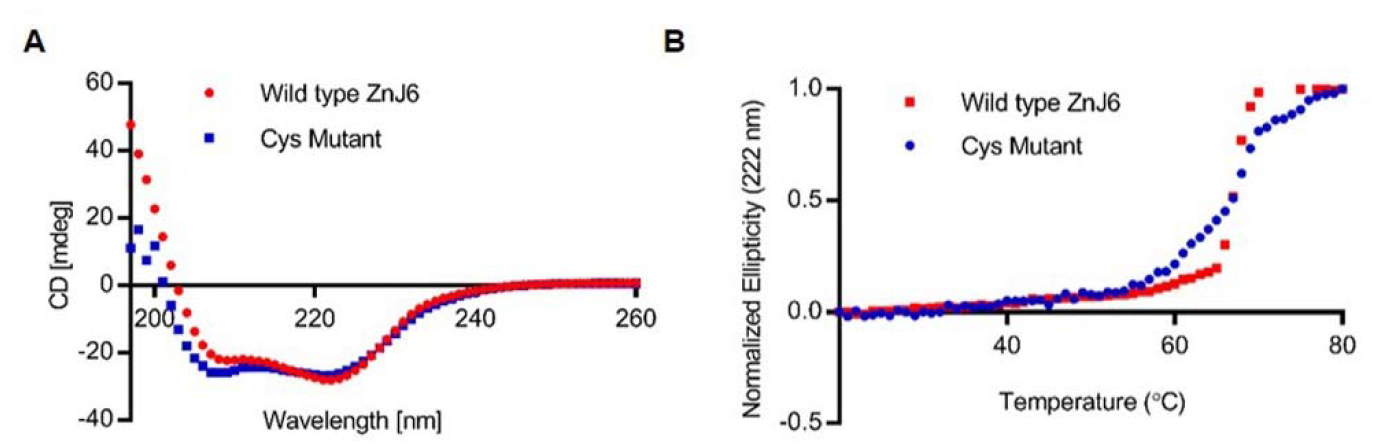
Recombinant ZnJ6 and cys-mutant are folded and stable up to 65°C. **A**, CD Single accumulation spectra ranging between 200-260nm was recorded for wild type ZnJ6 and its Cys-mutant at RT, to verify that the recombinant proteins are folded. **B**, The single wavelength melting curve was generated at a constant wavelength of 222nm at a temperature range of 20-80°C. Normalized CD melting curves of ZnJ6 and cys-mutant with relative values between 1.0 and 0.0 provide an overall comparative indication of the thermal stability of ZnJ6 and its cys-mutant. The wild type recombinant ZnJ6 shows a more coordinated structure at elevated temperatures, as indicated by the steep slope. However, both proteins have the same midpoint of the melting curve at 65°C.

### Recombinant ZnJ6 binds zinc through its cysteine-rich motif

To evaluate the zinc-binding ability of cys-rich domain, the Zn binding assay using 4-(2-pyridylazo) resorcinol (PAR) and *p*-chloro-mercurybenzoate (PCMB) was performed. The assay was done using 3μM of purified recombinant proteins (ZnJ6 and cys-mutant). The addition of 30 *µ*M PCMB caused the release of the zinc atom from the protein molecule, leading to the formation of a coloured complex due to its interaction with PAR, which was measured by absorbance at 500 nm (Hunt et al, 1985). Our results (Figure 3A and 3B) show that the Zinc finger domain of the protein is required for binding Zinc. Increased absorbance in the assay that contained ZnJ6 indicated the release of coordinated zinc from the protein by PCMB, which then formed a coloured complex with PAR. The recombinant protein purified from the bacteria was analysed just after purification (without pre-incubation) and after its incubation with ZnCl_2_ (with pre-incubation). Although zinc release was observed in both cases, there was relatively a lower release of the endogenous zinc ion if the protein was not pre-incubated with ZnCl_2_, since the recombinant protein purified from bacteria was not fully saturated with zinc. There was a negligible release of zinc ion from the cys-mutant, with or without pre-incubation, as it lost its coordination with zinc.

**Figure 3.**
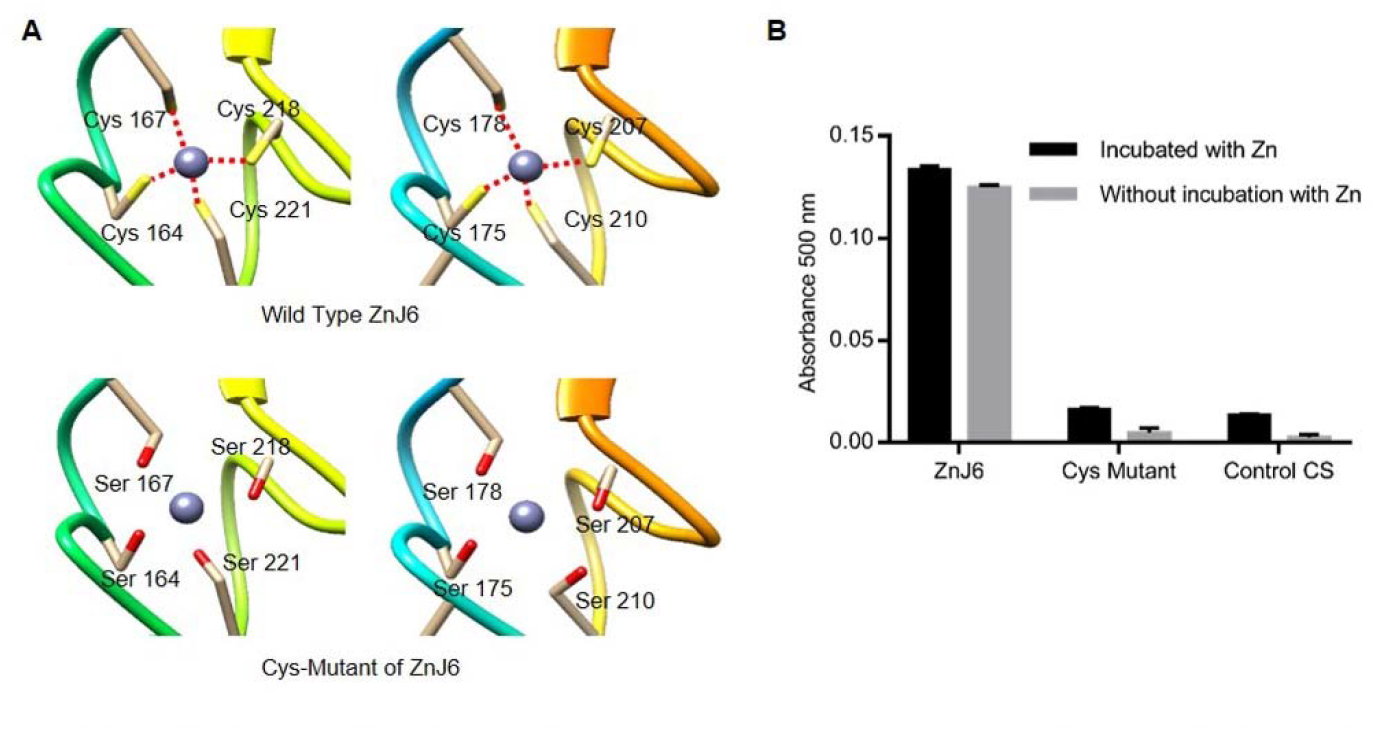
The cysteine-rich motif of ZnJ6 is required for its zinc-binding activity. **A**, The structure of the Zinc binding motif (wild type and cys-mutant) was predicted using homology modeling (constructed by UCSF Chimera). The predicted loss of coordination (red dotted lines) with zinc due to the replacement of cysteine with serine in the cys-mutant. **B**, Zinc binding activity of ZnJ6 (wild type and cys-mutant) was measured using 5 µM of the purified recombinant proteins, either pre-incubated with 40 µM ZnCl_2_ to completely saturate the protein with Zn, or monitored without pre-incubation with Zn, representing the Zn binding status of the protein extracted from bacteria. The release of zinc was obtained using para-chloromercuribenzoic acid (PCMB), and the PAR –Zn^+2^ complex that was formed was monitored at 500 nm. Citrate Synthase was used as a control.

### Recombinant ZnJ6 and its cys-mutant function as chaperones that prevent thermal aggregation of substrate protein Citrate Synthase

To establish the whether ZnJ6 can function as a chaperone, the classical Citrate synthase (CS) assay that measures the ability of the chaperone to prevent aggregation of a temperature-sensitive protein such as CS, was performed. This assay does not monitor refolding activities. CS is highly sensitive to temperature elevation and loses its folding already at 42°C, as shown in our controls and by Buchner et al. 1998. However, the CD measurements of ZnJ6 at increasing temperatures (Figure 2B) showed that it remained folded up to 65°C. The dose-dependent chaperone effect of ZnJ6 was determined by its incubation in increasing molar ratios relative to the CS substrate (ZnJ6: CS were 0.1: 1, 1:1, 2:1, 5:1, 10:1), at 42°C for 1 h. CS aggregation was measured by monitoring OD_360_ over time, whereby the increase in aggregation resulted in increased absorbance. The results indicate that ZnJ6 could function as a chaperone that prevented substrate aggregation, since, in the presence of ZnJ6, CS exposed to 42°C remained soluble even after 1 hour, starting at a ratio of 1:1 and reaching maximum protection of 86% when mixed with ZnJ6 in a 1:10 ratio. This protective activity (Figure 4), was dose-dependent. The requirement for the cysteine-rich domain in the chaperone activity was further examined by using the cys-mutant of ZnJ6. No significant difference could be recorded between the chaperone activity of the mutant and wild type ZnJ6 proteins, indicating that the ZnJ6 activity of preventing aggregation is independent of its cysteine-rich domain.

**Figure 4.**
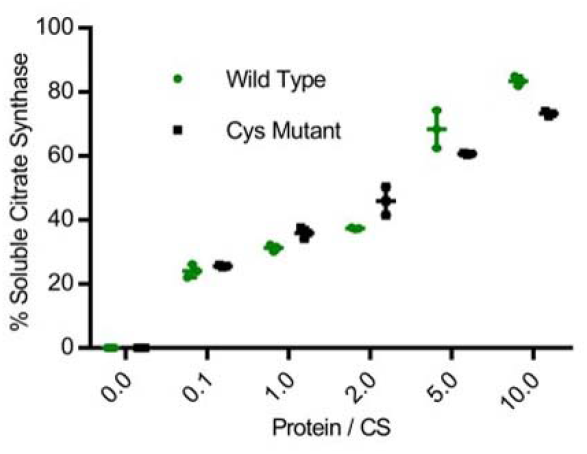
ZnJ6 and its cys-mutant have holding chaperone activity, measured by the Citrate Synthase thermal aggregation assay. Citrate synthase (CS) was diluted into a refolding mixture at 42 °C (1h) or into a refolding mixture with ZnJ6 (green circles). A parallel assay was also performed with the ZnJ6 cys-mutant (black squares). All assays were performed in the presence of increasing molar ratios as compared to CS (ZnJ6: CS; 0:1, 0.1: 1, 1:1, 2:1, 5:1, 10:1). The thermal aggregation of CS was measured by monitoring OD_360_ over time. The absorbance was used to calculate the percentage of soluble CS after 1 h of incubation at elevated temperatures. CS that was not subjected to increased temperature served as the 100 % soluble control. CS alone incubated at 42°C for 1 h served as the fully aggregated substrate.

### ZnJ6 is unable to reduce the disulfide bonds of insulin but prevents its aggregation in a reducing environment

The thiol-dependent activity of the protein was examined in the insulin turbidity assay (Arne, 1979). Insulin contains two polypeptide chains, α and β, that are held together by disulfide bridges. In this assay, a reducing agent such as DTT is added to the mixture, causing the disulfide bridges that hold the two polypeptide chains together to open. Once reduced and released, the β -subunit of insulin aggregates and its precipitation can be measured at 650 nm. In contrast to thioredoxins that accelerate precipitation of the insulin β chain, ZnJ6 did not have such a reducing power. However, when insulin was reduced in the presence of DTT, the addition of ZnJ6 in increasing molar concentrations relative to insulin (CS: Insulin 0:1, 0.2:1, 0.5:1 and 1:1) prevented the β chain precipitation in a dose-dependent manner (Figure 5A). Furthermore, unlike the wild-type protein, the cys-mutant failed to prevent precipitation of the insoluble reduced insulin chain with the same efficiency (Figure 5B-C). Thus, although ZnJ6 lacked any reducing activity, it could prevent insulin chain precipitation, and the zinc-binding motif appeared to have a role during the prevention of aggregation. To further confirm the role of the cysteines in preventing the aggregation of insulin chains by ZnJ6, we monitored the amount of - SH groups with and without the Insulin substrate, using 5-dithio-bis-(2-nitrobenzoic acid) (DTNB), also known as the Ellman’s reagent. DTNB binds to reduce -SH groups, forming a coloured complex that can be measured by its absorbance at 412 nm. We expected that prevention of insulin aggregation occurred when the ZnJ6 -SH groups were occupied by the insulin -SH groups forming disulfide bridges, thereby decreasing the total amount of -SH groups in the solution. The amount of protein-bound SH (PB-SH) groups in the mixture was determined before and after incubation (2 h) of recombinant ZnJ6, with and without insulin, in the presence of 1 mM DTT. To calculate the PB-SH, first DTNB was added to the mixture, and total –SH groups (T-SH) were measured by taking absorbance at 412 nm, [before precipitation with Trichloroacetic acid (TCA)]. Next, a parallel mixture was TCA precipitated, centrifuged to remove the proteins, and DTNB was added to the protein-free supernatant. This step eliminated the effect of remaining DTT on binding to DTNB and the value obtained was NP-SH. The difference between the T-SH and the NP-SH values gave the protein bound SH (PB-SH) groups that was calculated before and after the 2h incubation of the different mixtures. Our results, shown in Figure 5, indicate that the SH groups of ZnJ6 remained unchanged when ZnJ6 was incubated alone, along with DTT. However, when ZnJ6 was incubated in the presence of Insulin in an equal molar ratio, the amount of protein bound SH groups decreased dramatically after the incubation with Insulin, thus supporting the formation of disulfide bridges between ZnJ6 and the reduced Insulin chains (Figure 5D). In conclusion, ZnJ6 lacked any reducing activity by itself, as it failed to precipitate the Insulin chain in the absence of DTT. However, it did prevent the aggregation of reduced Insulin chains by forming disulfide bridges between the cys-rich motif of wild type ZnJ6 and reduced insulin chain.

**Figure 5.**
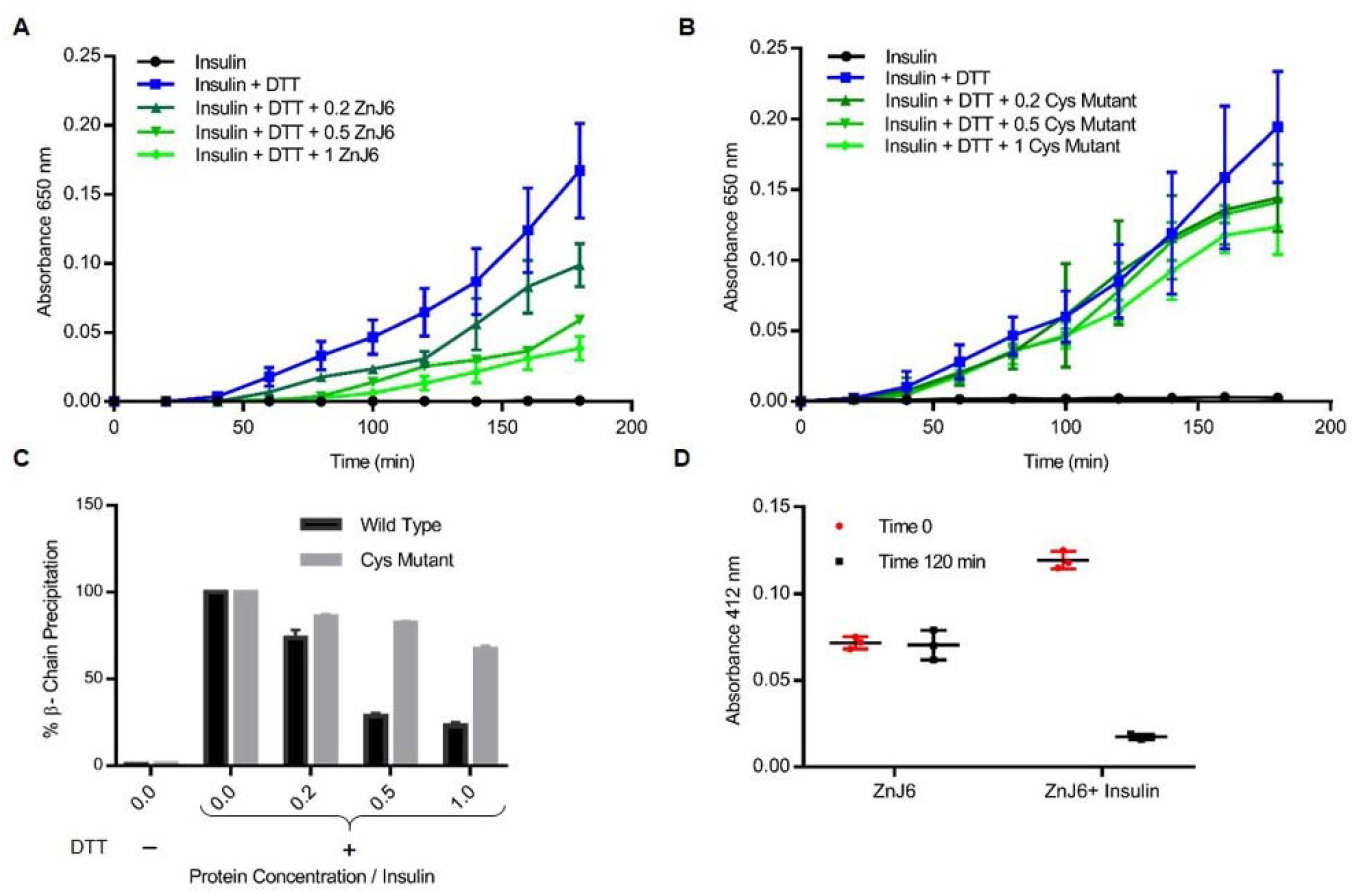
The Zinc finger domain is required for preventing aggregation of the reduced insulin chain. The Insulin turbidity assay was used to examine the thiol-dependent activity of ZnJ6. Increasing molar ratios of ZnJ6 as compared to insulin were added (0:1, 0.2:1, 0.5:1, 1:1), and precipitation of the insulin β-chain was measured at 650 nm, during 3 h at 25°C. A reaction containing insulin alone served as control. **A**, Precipitation was monitored in the presence of wild type recombinant ZnJ6 and **B**, its cys-mutant. **C**, Summary of end-point precipitation values obtained in the presence of different molar ratios of the ZnJ6 (black columns) and its Cys-mutant (grey columns) after 3 h. **D**, Protein-bound sulfhydryl (-SH) groups were calculated using the Ellman’s test, before (red circles) and after incubation of ZnJ6 (black squares), with or without insulin, that was introduced in an equal molar ratio.

### ZnJ6 promotes the oxidative refolding of RNase A

To further investigate the thiol dependent oxidative refolding ability of ZnJ6, the oxidative refolding of reduced and denatured RNaseA (rdRNaseA) was measured in the presence of increasing molar ratios of ZnJ6. The native structure of RNaseA is stabilized by four disulfide bridges. Once these bonds are reduced, and the protein is denatured by guanidinium HCl, it loses its activity. Recovery of rdRNaseA enzymatic activity requires the native disulfide bonds to reform and to stabilize the refolded structure. Upon removal of the guanidinium HCl denaturant, the activity of rdRNaseA alone failed to regain its activity rapidly by spontaneous refolding. The reason for this failure could originate from the formation of non-native disulfide bridges and for the relatively long time required for proper refolding. However, refolding and reactivation occurred in the presence of ZnJ6 in increasing molar ratios to rdRNaseA (0:1, 0.2:1, 0.6:1 and 1.2:1). The activity of rdRNaseA was restored in a gradual manner, starting from 20% when added in a 0.2:1 molar ratio and reaching up to 50% within an hour following removal of the denaturant, when ZnJ6 was added in 1.2 molar ratio to rdRNaseA, (Figure 6). Similar molar ratios of the cys-mutant showed only a minimal effect and failed to refold the RNaseA with the same efficiency. These findings indicate that ZnJ6 could assist the reformation of native disulfide bridges in the reduced and denatured RNase A, thus regaining its activity.

**Figure 6.**
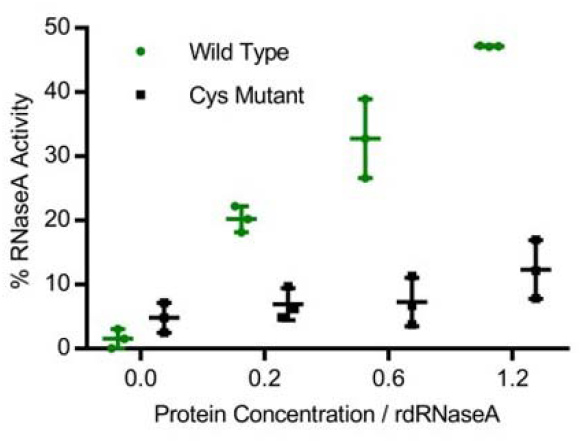
ZnJ6 affects the refolding of reduced and denatured RNaseA. Refolding of reduced and denatured RNaseA (rdRNaseA, generated using GnHCl and DTT) was initiated by its 200-fold dilution into renaturation buffer to a final concentration of 25μg/mL. Reactivation was conducted with and without ZnJ6, which was added in molar ratios of 0:1, 0.2:1, 0.6:1, and 1.2:1 as compared to rdRNaseA. Reactivation was monitored in the presence of the WT ZnJ6 (green circles), or its cys-mutant (black squares). Aliquots were removed at various intervals and transferred into the assay mixture, and RNaseA activity was measured by monitoring the hydrolysis of cytidine 2’3’-cyclic monophosphate at 284 nm. The points represent the final percentage of RNaseA activity as compared to native RNaseA, after 1 h of refolding.

### ZnJ6 interacts with photosynthetic proteins

The protein interactome plays an important role in predicting the function of target proteins. We, therefore, performed pull-down assays to examine the interacting partners of ZnJ6 that served as a bait for these assays. Since we encountered difficulties in overexpressing the SBP-tagged ZnJ6 *in vivo* to a level that could support efficient affinity purification, we carried out the experiment using the recombinant protein. Recombinant SBP-tagged ZnJ6 (100 μl, 10μM) was bound to a streptavidin column and further incubated with *C. reinhardtii* chloroplast extracts (4 ml, 0.1 μg/μl) for 2 hours. The beads were washed to remove the non-specific proteins until reaching a protein-free wash fraction (usually after washing with five column volumes). Finally, the recombinant ZnJ6-SBP bait protein was eluted with biotin, along with its interacting proteins (Supplemental Figure S4). Recombinant SBP-tagged MBP was used as an experimental control that was treated similarly. The eluted fractions were analyzed by mass spectrometry (MS). MS data were analysed using the MaxQuant software. Protein identification was set at less than a 1% false discovery rate. Label-free quantification (LFQ) intensities were compared among the three SBP-ZnJ6 biological repeats and the three SBP-MBP repeats with the Perseus software platform using the student t-test analysis.

The enrichment threshold (LFQ intensities of SBP-ZnJ6 subtracted from the SBP-MBP controls) was set to a log2-fold change ≤ -3 (8 fold enrichment as compared to control) with p< 0.05. The filtered proteins were categorized to functional groups both manually and by BLAST2GO, based on enrichment for Biological Processes. The manual categorization suggested that the ZnJ6 interacting proteins comprised of photosynthetic PSI proteins, transporters, ubiquinols, chaperones, and chlorophyll-binding proteins (Figure 7A, 7B, and Supplemental Table 1). It is riveting that ZnJ6, unlike maize BSD2, did not interact with the highly abundant subunits of Rubisco (LS/SS) (Salesse et al., 2017)(Li et al., 2020), supporting the binding specificity between ZnJ6 and its associated proteins. A broader classification was done using BLAST2GO enrichment analysis, setting a minimum threshold of two-fold enrichment for the associated proteins, as compared to their gene abundance in the genome data set. Using this approach, we also observed the high enrichment of photosynthetic proteins in the ZnJ6 interactome, along with proteins that related to oxireductases, metabolic enzymes and.transporters. The association with proteins of photosynthetic complexes supports the possibility that ZnJ6 could have a targeted role in the assembly of such photosynthetic complexes.

**Figure 7.**
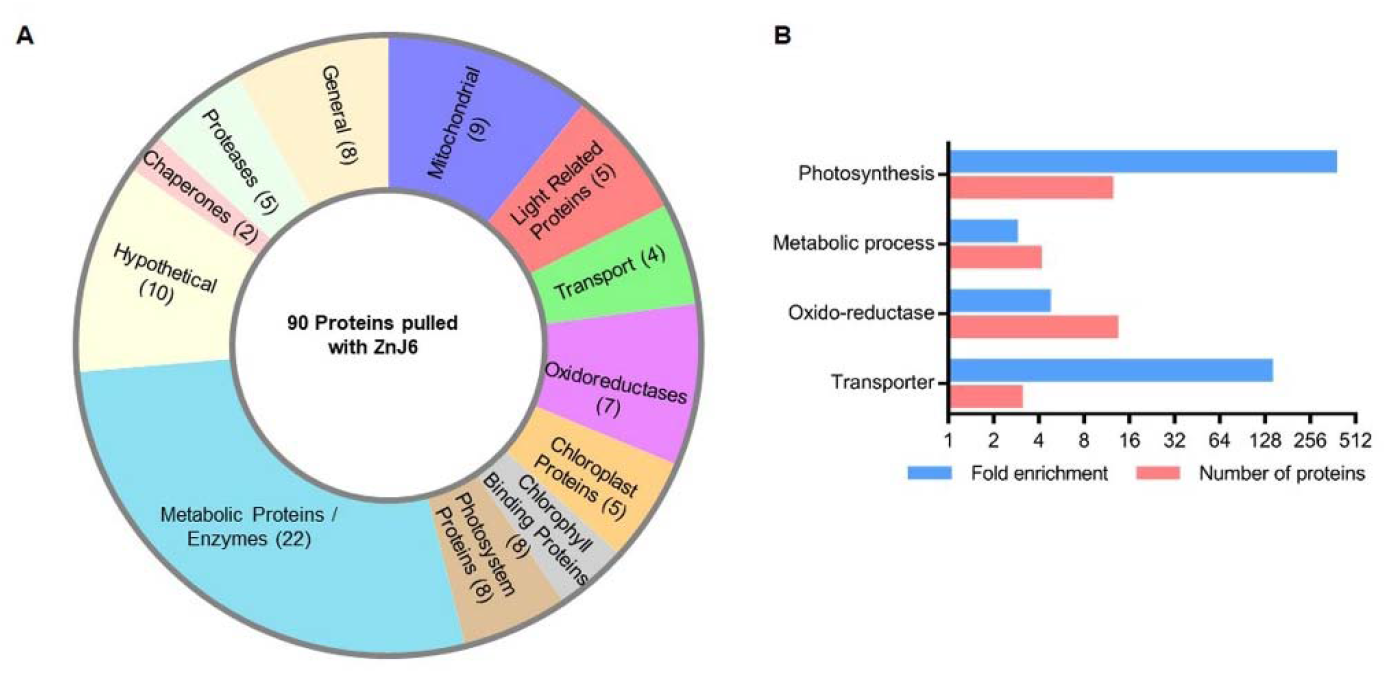
Protein categories that are associated with ZnJ6 in pull-down assays. The proteins pulled down with ZnJ6 were determined by LC-MS/MS analysis, in triplicates, and compared to control pull-down assays performed with a non-related MBP protein that was treated similarly. The proteins were identified by the MaxQuant software using Phytozome database annotations. Differences between the proteomic contents of the ZnJ6 and MBP pulled-down fractions were determined using the Perseus statistical tool. Proteins with eight-fold enrichment as compared to the control, with p< 0.05, were categorized. **A**, Manual categorization of proteins into functional groups along with their relative abundance, represented by the respective area and number of proteins (shown in brackets). The hypothetical group contains proteins with non-defined functions. **B**, BLAST2GO enrichment determined by the biological process, setting a minimum threshold of two-fold enrichment for the associated proteins as compared to their gene abundance in the genome data set.

### *Chlamydomonas* mutant cells expressing a low level of ZnJ6 (ΔZnJ6) are more tolerant to oxidative stress but sensitive to reductive stress

As ZnJ6 is a redox-regulated chaperone, we wanted to understand how it affects the cells in changing redox environments. For this, a *Chlamydomonas* insertional mutant expressing a low level of ZnJ6 (CLiP mutant, LMJ.RY0402.048147) denoted ΔZnJ6, was examined. The ΔZnJ6 mutant was first confirmed using colony PCR and western analysis, using anti-ZnJ6 antibodies (Supplemental Figure S5). To examine the effect of this mutation on growth and resistance to different oxidizing environments, cells were grown to mid-log phase in High Salt (HS) medium, with a light intensity of 150 µmol/s/m^2^, in the presence of paromomycin, for selective maintenance of the mutation. The cells were exposed to different concentrations of H_2_O_2_ (0, 2, 5, 10 and 20 mM) and MeV (0, 2, 5, 10 and 20 µM) for an hour. Cells were then washed, spotted and allowed to grow on HS plates for 5 days at 23°C. We observed that under increased oxidizing conditions, the mutants appeared to be more tolerant to oxidative stress than the wild type cells. However, the wild type and ΔZnJ6 cells presented insignificant variations in their growth under optimum or mild oxidizing conditions (up to 2 mM of H_2_O_2_ and 2 µM of MeV), Figure 8A and 8B.

**Figure 8.**
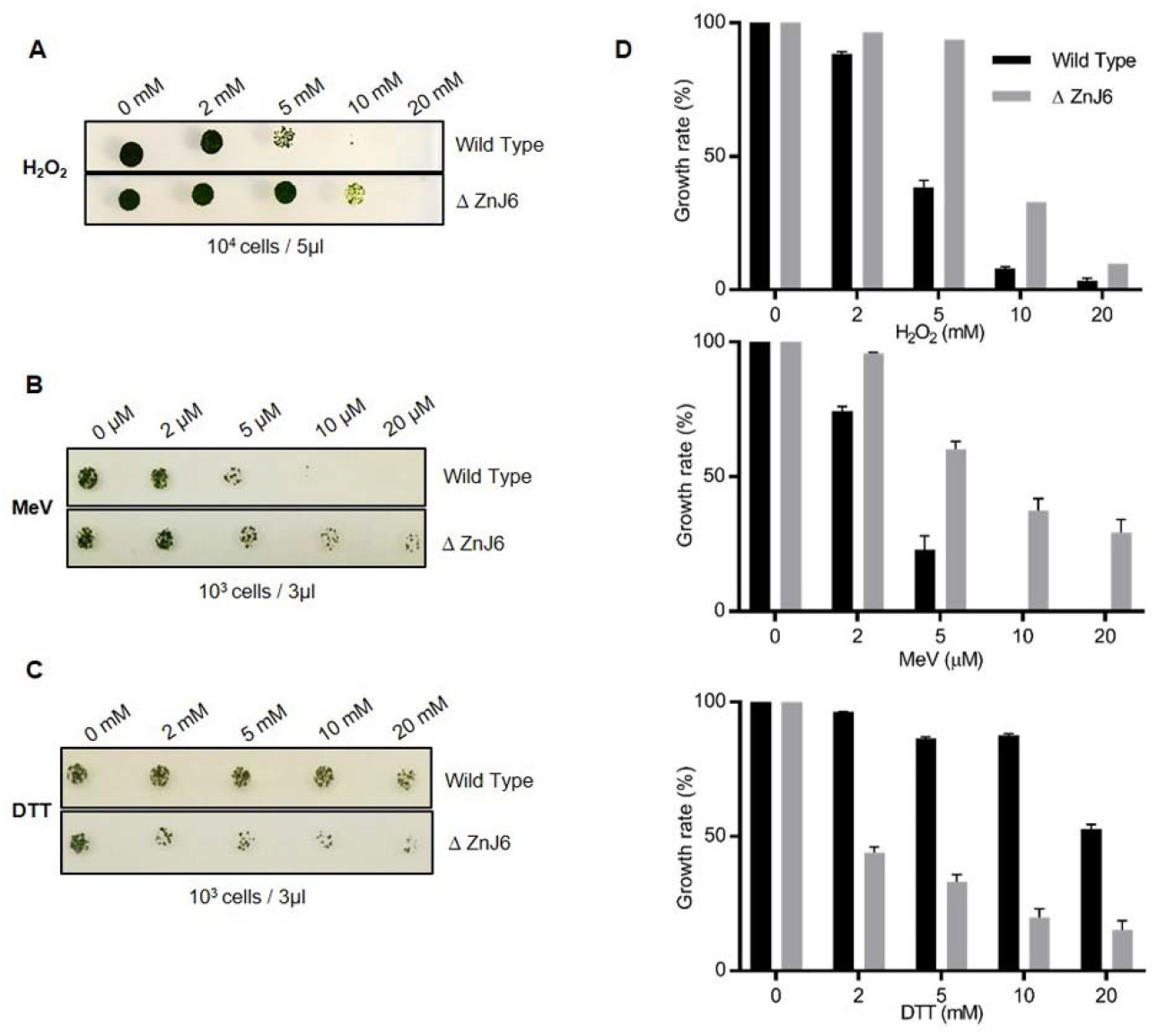
ZnJ6 knock-down mutant of *C.reinhardtii* (ΔZnJ6) shows tolerance to oxidative stress but increased sensitivity to reducing conditions. **A, B**, Wild type, and ΔZnJ6 cells were exposed to oxidizing conditions by incubation with increasing concentrations of H_2_O_2_ and MeV for 1 hr. The cells were washed, and growth was monitored on HS plates. **C**, Cells were exposed to reducing conditions by incubation with DTT for 1 hr. The cells were washed, and growth was monitored by plating over HS plates. **D**, Growth variations of cells treated with H_2_O_2_, MeV, or DTT, as shown in panels A-C were measured using the Multi-gauge software. The growth of untreated cells served as control (100%).

An opposite effect was observed when the growth of the mutant cells was compared to wild type cells under reducing conditions. Cells were treated with increasing concentrations of DTT (0, 2, 5, 10, and 20 mM) for 2 hours, washed and spotted on HS plates, and allowed to grow for 5 days at 23°C. While wild type cells grew well in the presence of all DTT concentrations, growth of the mutant cells was severely compromised (Figure 8C). Growth under reducing conditions can possibly lead to structural changes of protein, resulting in their toxic aggregation. Since the growth of the ?ZnJ6 mutant was impaired in the presence of DTT as compare do wild type cells, we assume that ZnJ6 could be involved in preventing the massive reduction caused by DTT *in vivo*, thereby making the wild type resistant to the DTT induced stress. This hypothesis also corroborates with our finding that ZnJ6 could prevent the aggregation of reduced insulin chain in vitro.

### The ZnJ6 insertional mutant is sensitive to heat stress

To further elaborate on the role of ZnJ6 during exposure of the cells to other stress conditions, the response to a commonly encountered mild heat stress was evaluated in ΔZnJ6. In this assay, we spotted 5 µl of cells with increasing cell concentrations on HS plates and allowed them to grow for different time periods at 37°C (0, 1, 2, 4 h). Following this treatment, the plates were transferred back to 23°C and the cells were allowed to grow for five additional days. Figure 9A shows that exposure to the increased temperature for 2 h or more resulted in chlorophyll reduction, since the cells became yellow. Also, growth of the mutant cells was slower as compared to the wild type control. Further on, a loopful of cells taken from spots seeded with 10^5^ cells in each treatment were resuspended in 100 µl HS media, and 5 µl of cells were spotted on the new HS plate and allowed to grow for additional 5 days at 23°C. In this case too, the damaging effect of the temperature stress on the growth of the mutant cells continued, as growth was still impaired (Figure 9B). Image analysis, using myImageAnalysis software, also confirmed this observation. Growth of both wild type and ΔZnJ6 cells was reduced in response to the heat stress, as compared to their growth at 23°C, but the mutant had a significantly impaired growth as compared to wild type cells (Figure 9C, D). We concluded that ZnJ6 might assist the cells in withstanding heat stress, thus allowing continued growth.

**Figure 9.**
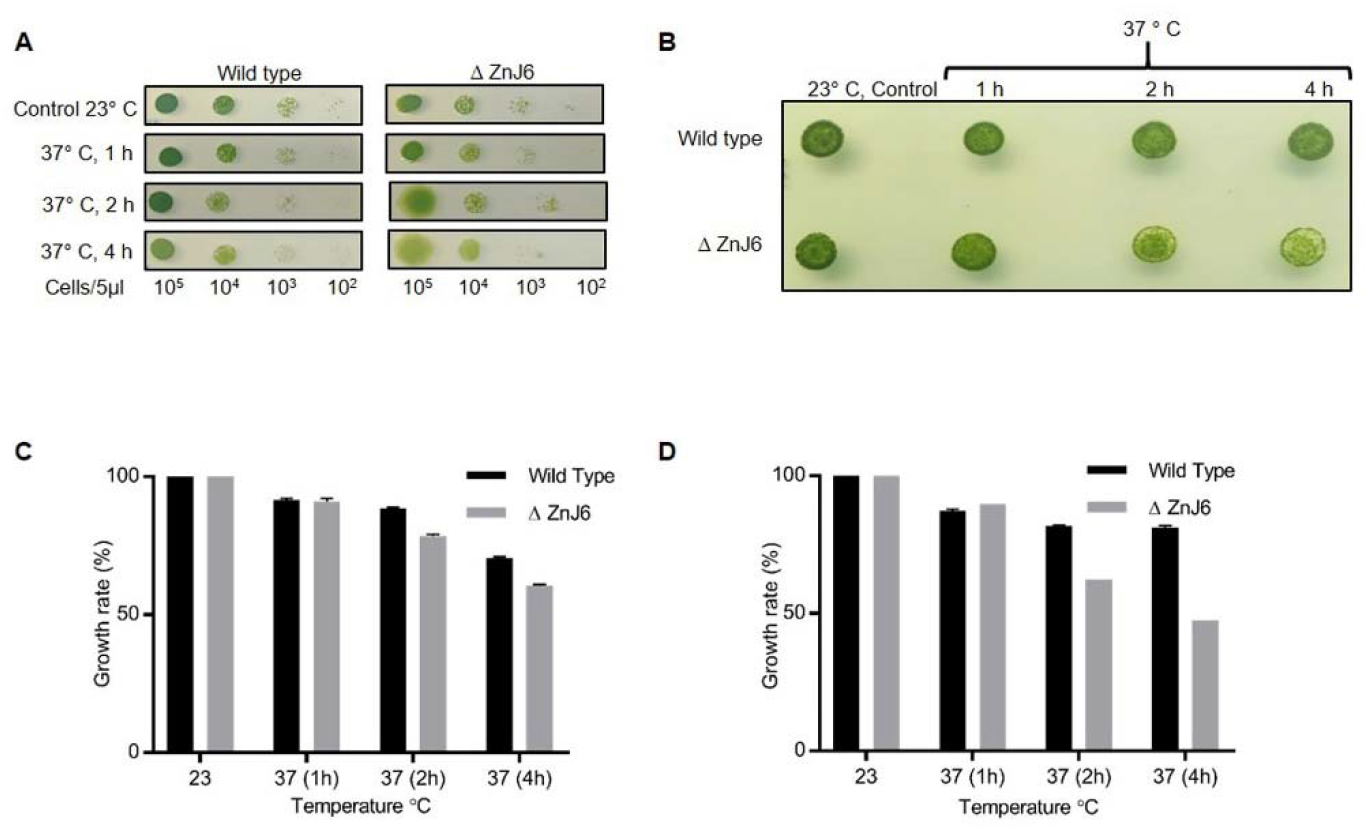
ZnJ6 knock-down mutant (ΔZnJ6) is sensitive to temperature stress. **A**, *Chlamydomonas* cells (wild type and ΔZnJ6) grown to mid-log phase in HS medium, were serially diluted. Aliquots (5μl) from each dilution were spotted on HS plates and incubated at 37°C for 1, 2, and 4 h. Control plates were maintained at 23°C. All the plates were then allowed to grow for 5 days under continuous illumination at 23°C. Decreased growth and an increase in yellowing of the cells is observed in ΔZnJ6 cells that were incubated at increased durations of heat treatments. **B**, The reduced growth of the ΔZnJ6 cells is maintained even in a continued growth when the cells from Panel A were further sub-cultured and replated. After 5 days of growth, a loopful of cells (1 µl loop taken from the cells spotted at a dilution of 10^5^ cells per 5 µl) was resuspended in the HS media, spotted, and allowed to grow for additional 5 days at 23°C. Mutant ?ZnJ6 cells show reduced growth when treated at 37°C for 2-4 h. **C and D**, Densitometric analysis of the growth observed in panels A (taken from cells spotted at a dilution of 10^3^ cells per 5 µl) and B, respectively, were measured using myImageAnalysis (Thermo Fisher) software. The histograms were plotted after subtracting the background intensity from the measured intensity and considering the control as 100%.

## DISCUSSION

The *C. reinhardtii* thylakoid membrane-associated DnaJ-like protein ZnJ6 (Cre06.g251716), is a zinc finger oxidase that contains four cysteine-rich CXXCX(G)X(G) domains. These form two C_4_ type zinc fingers, responsible for binding Zinc. The Zinc binding domain provides a more stable coordinated structure to the protein at elevated temperatures, as suggested by circular dichroism. Domain prediction by the TMHMM and Phobius servers (Supplemental Figure S2) identified a transmembrane domain in ZnJ6, thus excluding the possibility that it could serve as a potential ortholog of BSD2 from higher plants (GRMZM2G062788_T01) that does not have any evidence for being a transmembrane protein. This is also supported by the association of ZnJ6 with the thylakoid membrane fraction using biochemical fractionation assays. Thus, ZnJ6 appears to be a distinct protein.

Zinc finger domains are known to be involved in protein-protein interactions and can even contribute to the ability of chaperones to identify substrate proteins in their denatured state (Szabo et al, 1996). To understand the role of the cysteine-rich motif in chaperone activities of ZnJ6, wild type, and cys-mutant recombinant proteins were analyzed using classical *in vitro* chaperone assays. Based on the Citrate Synthase prevention of aggregation assay, ZnJ6 is shown to have a chaperone “holding activity” regardless of the mutation in the cysteine-rich domain. However, this domain was required for the redox activity of this protein. For example, ZnJ6 failed to induce precipitation of the Insulin β-chain in the insulin turbidity assay, and this activity required the cys-rich domain. The function of the Zn-binding domain in ZnJ6 varied from that of thioredoxin (Arne, 1979) (Jeon and Ishikawa, 2002), since the latter could induce precipitation of the Insulin β-chains, whereas ZnJ6 could only protect these chains from precipitation, in the presence of a reducing agent such as DTT. We, therefore, concluded that ZnJ6 lacks independent reducing activity of disulphide bonds in its target proteins. Overall, this also excluded the possibility that ZnJ6 possessed protein disulphide isomerization activity (PDI). Based on the RNaseA assay, we showed that ZnJ6 has an oxidizing activity, and it assists in the native folding of reduced-denatured RNaseA (rd RNaseA). This activity was dependent on the presence of a functional Zn-binding domain. We concluded that ZnJ6 is a chaperone that can hold its target to prevent aggregation; it lacks reducing activity but can promote the disulphide-bridge formation in its target proteins.

Sub-cellular localization and protein interaction studies are fundamental to elucidate the mechanism and context of protein functioning (Goodin, 2018). Thus, using subcellular fractionation analysis, we identified ZnJ6 in the thylakoid membrane. This finding was found to be in agreement with the bioinformatically predicted transmembrane domain in ZnJ6. To further expand our general understanding of its context and protein interactions, we performed a pull-down assay in which we affinity-purified chloroplast extracts of *Chlamydomonas* over immobilized recombinant ZnJ6. This approach showed that ZnJ6 co-purified with the majority of photosynthetic proteins (12), oxidoreductases (13), proteases (5), and chaperones (2), suggesting that ZnJ6 could be responsible for chaperoning a multitude of substrate proteins. However, at this stage, we still cannot relate these findings to a direct interaction between these proteins and ZnJ6. The association of ZnJ6 with oxidoreductases could indicate its involvement in maintaining a subcellular redox balance, while its co-purification with photosynthetic proteins (with the majority of photosystem I proteins) could indicate its role during the assembly of the photosynthetic complex.

We have observed the enrichment of metabolic enzymes in the BLAST2GO analysis. This observation is in agreement with growing evidence for dual functions observed for metabolic enzymes, among them RNA binding activities. Such activities could be related to their moonlighting activities, yet other explanations are offered. We previously reported that Rubisco LSU possesses RNA binding activity, which was related to its regulation under oxidizing conditions (Yosef et al., 2004). The RNA-binding activity of metabolic enzymes was recently expanded to a multitude of enzymes (Perez-Perri et al., 2018), raising interesting possibilities for such activity (Sachdeva et al., 2014).

To elaborate on the role of ZnJ6 in redox responses, we performed in vivo experiments monitoring the growth of the *C. reinhardtii* ZnJ6 knock-down mutant cells that expressed a low level of ZnJ6 in the different redox environment. These assays indicated that the ?ZNJ6 mutant cells were more tolerant to oxidative stress caused by short incubations with H_2_O_2_ or MeV, as compared to wild type cells. However, the mechanism behind this activity is still unclear. A possible explanation could be that in the absence of ZnJ6, the glutathione pool was shifted to its reduced form, thus enabling a damaging reduction of target proteins that prevented their function. A similar physiological effect was reported for the FtsH5-Interacting Protein (FIP, At5g02160) in *Arabidopsis*. This protein is reported only in mosses and higher plants and is another DnaJ-like protein that lacks the typical J-domains (Lopes et al, 2018). ZnJ6 and FIP have four and two cysteine-rich motifs, respectively; and mutants of both proteins show tolerance to oxidative stress. ZnJ6 associates with an FtsH-like protease, as well as with members of the ClpP protease complex. ClpP proteases are involved in the maintenance of chloroplast protein homeostasis (Adam et al., 2006) (Nishimura and Van Wijk, 2015). Also, chloroplast chaperones are known to regulate protease activities and function in synergy to maintain protein quality control (Nishimura et al, 2017). Therefore ZnJ6 could be involved in protein quality control and in regulating protein homeostasis.

In contrast to the improved growth of the ΔZNJ6 mutant cells under oxidizing conditions, these cells were more sensitive to a reducing force induced by DTT, showing impaired growth as compared to wild type cells. We assume that in the presence of DTT, protein structures may be affected due to the reduction of disulfide bridges, as these are required to stabilize proteins, among them also those involved in photosynthesis. Thus, the oxidizing activity of ZnJ6 could recover disulfide bridge formation, thus protecting protein structures in the wild type cells. In its absence, the reductive force of DTT could impair protein structures, possibly also leading to their aggregation. Thereby, ZnJ6 could provide resistance to the wild type cells against DTT induced stress. The in vitro insulin aggregation assay also supports the observation.

It was earlier shown that shifting *Chlamydomonas* cells from 25°C to 37°C induced a heat stress response (HSR) (Schroda et al., 2015). Our data suggest that ZnJ6 is involved in protection against a heat stress, since exposure of the ?ZNJ6 mutant cells to elevated temperatures for 2-4 hours resulted in degradation of chlorophyll and impaired growth. Thus the mutant cells appeared to be sensitive to heat stress (37°C) much more than the wild type cells, possibly highlighting the importance of the chaperone activity of ZnJ6 under a temperature stress. This is further supported by the finding that ZnJ6 was found to interact with ClpP proteases that are involved in chloroplast unfolded protein response, UPR and in proteostasis processes (Ramundo et al., 2014). The ability of ZnJ6 to protect a substrate protein from heat induced aggregation was shown *in vitro* in the CS assays. Its structure is also stable up to 65°C, enabling such activity. ZnJ6 was also shown to interact with the temperature-sensitive catalytic chaperone Rubisco activase in the pull-down assay. This could suggest that it has a role in the prevention of irreversible aggregation of temperature-sensitive proteins in the chloroplast.

## CONCLUSION

Here we show that ZnJ6 is a chaperone that in addition to its ability to prevent aggregation of misfolded substrate proteins, possesses an oxidizing activity that can restore reduced disulfide bridges, thus stabilizing protein structure. We therefore suggest that ZnJ6 assists in maintaining the redox balance in the chloroplast. ZnJ6 is a thylakoid membrane protein, shown to interact with photosystem complexes. As shown for other DnaJ-like proteins such as PSA2 (Fristedt et al., 2014) and LQY1 (Lu et al., 2011), ZnJ6 could also be involved in similar assembly processes, although its localization in the thylakoid membranes could restrict its function to complexes that are formed in the membranes. Alternatively, ZnJ6 could also affect the biosynthesis of thylakoid membrane components, as these are also coordinated with the photosynthetic machinery (Bohne et al., 2013). Further studies are required to deepen our understanding of ZnJ6 function and role during different stresses.

## MATERIAL AND METHODS

### Isolation of RNA, cDNA synthesis and cloning

Early log-phase cells (OD_750_ = 0.25-0.35, 2×10^6^ cells/ml) were used to isolate total RNA by using TRI Reagent (Sigma) protocol. The cDNA was synthesized using High capacity cDNA reverse transcription protocol (Applied Biosystems) with 1 µg RNA as a template. Bacterial clones for recombinant protein expression were generated as described in the supplemental methods section and the primers used were mentioned in supplementale tables 2 and 3.

### *Chlamydomonas* strains and Growth Conditions

*Chlamydomonas* strains (cc-125 and cc-4533) were grown and maintained on TAP plates at 23°C. The knock-down ZnJ6 CLiP mutant LMJ.RY0402.048147 (ΔZnJ6) was obtained from the *Chlamydomonas* Resource Center (Li et al, 2019). The knock-down mutant was maintained over 10 µg/ml Paramomycin in Tris-acetate-phosphate (TAP) plates and verified by colony PCR followed by western analysis. The fresh colony was first inoculated in 10ml TAP media with required antibiotics followed by large scale culturing in High Salt (HS) or TAP media (as per requirement), with 12 h dark/light cycles (at 150 µmol/s/m^2^) and constant rotary shaking at 100 rpm.

### Recombinant protein purification

An overnoght bacterial stater culture (10 ml) in LB medium supplemented with 100 μg/ml ampicillin and 25 μg/ml chloramphenicol (for rosetta strain only) was inoculated into 1 L of LB, supplemented with required antibiotics and 1% glucose. Expression of the SBP-tagged pMBP-GB1-ZnJ6 (see supplemental methods) was induced upon the addition of 0.2 mM IPTG when cells reached OD_600_ = 0.5–0.7, at 20°C for 16 h. The culture was harvested and resuspended in lysis buffer (20 mM Tris-HCl, pH 7.4, 200 mM NaCl, 1 mM EDTA) containing 0.1% Brij 58, Sigma Aldrich, a protease inhibitor (PI) cocktail, and 5 μg/ml DNaseI. The cells were disrupted in a French Press at 1500 psi and centrifuged at 45,000 rpm (Beckman 70 Ti rotor). The supernatant was loaded onto an amylose column (NEB). After washing the column with 5 column volumes of lysis buffer, ZnJ6 was eluted with 10 mM maltose in the same buffer. Next, the protein was cleaved using the TEV protease, to remove the MBP tag. The SBP tagged cleaved protein was purified again over the streptavidin-Sepharose (A2S) column. Protein concentration was estimated using the BCA protein assay kit (Thermo scientific). The the SBP-tagged cys-mutant and MBP proteins were purified similarly. Elution fractions were analysed on 15% SDS-PAGE (Supplemental figure S3).

### Subcellular fractionation of Cytoplasmic and chloroplast fractions

Mid-log cells (1L) were harvested and disrupted by nitrogen cavitation in a Yeda Press Cell Disruptor at approximately 100 PSI. The disrupted cells were centrifuged at 2000g in Corex glass tubes. The supernatant contained the cytoplasmic fraction. The pellets were resuspended in 6 ml of Percoll buffer (330 mM Sorbitol, 1 mM MgCl_2_, 20 mM NaCl, 2 mM EDTA, 1 mM MnCl_2_, 2 mM NaNO_3_, 5 mM Na-ascorbate and 50 mM HEPES, pH 7.6). A sample of 5ml was loaded over a 45/70% Percoll step gradient, which was centrifuged at 20,000g for 10 min at 4°C ina SW40 rotor. The intact chloroplasts were collected from the interphase of the step gradient. The efficiency of the chloroplast isolation was determined by measuring the chlorophyll content in the purified fractions.

The subcellular fractions were verified by western blot analysis using gels loaded with equal protein quantities. Antibodies against organelle-specific proteins were used to verify the subcellular fractions. These targeted the Rubisco small subunit (rbcS) and the 33-kDa oxygen-evolving enzyme (OEE33)as chloroplast markers, HSP70A as a cytoplasmic marker. Antibodies against the 32 kDa psbA which encodes for the D1 protein of photosystem II served as a marker for the chloroplast and its thylakoids. Antibodies against the recombinant ZnJ6 fragment 1-165 are described in Supplemental Figure S1.

### Separation of the membrane and soluble protein fraction

*Chlamydomonas* cells (250 ml) were grown, pelleted as above, and resuspended in 10 ml, 25 mM HEPES-KOH, pH 7.5, 5 mM MgCl_2_, 0.3 M (10.2%) sucrose with PI. The resuspended pellet was then disrupted with the Yeda Press apparatus at 500 PSI. Membrane and soluble fractions were separated by centrifugation at 100,000 g for 1 h at 4°C. The soluble proteins were precipitated using 20% TCA (final concentration) for 1h at 4°C and washed twice with 100% acetone. Pellets of both soluble and membrane proteins were dissolved in 40 mM Tris-HCl pH 7.4, 5 Mm EDTA, 4% SDS (Wittkopp et al, 2018).

### Isolation of Thylakoid membranes

Thylakoid membranes were isolated by harvesting a 250 ml culture of mid-log cells grown as described above. Cells were pelleted at 4500g for 10min at 4°C and resuspended in 25 mM HEPES-KOH, pH 7.5, 5 mM MgCl2, and 0.3 M sucrose supplemented with a cocktail of a PIs. Cells were disrupted using nitrogen cavitation in the Yeda Press Cell Disruptor at 500 PSI and centrifuged at 2,316g (10 min, 4°C), to separate between the membrane and soluble fractions. The pellet was resuspended in 5 mM HEPES-KOH, pH 7.5, 10 mM EDTA, 0.3 M sucrose, and a cocktail of PIs, followed by centrifugation at 68,600 g for 20min at 4°C. The resulting pellet was resuspended in 5ml, 5 mM HEPES-KOH, pH 7.5, 10 mM EDTA, 1.8 M sucrose, and a mix of protease inhibitors. The resuspended sample (at the bottom of the tube) was carefully overlaid with 2 ml 5 mM HEPES-KOH, pH 7.5, 1.3 M sucrose, 10 mM EDTA, and then 5 ml, 5 mM HEPES-KOH, pH 7.5, 0.5 M sucrose. The thylakoid membranes were then centrifuged at 247,605 g for 1 h at 4°C. The Thylakoid membranes were collected from the interface between the fractions containing 1.8 M and 1.3 M sucrose in the above-mentioned gradient. The thylakoid membranes were spun down at 68,600 g for 20 min at 4°C and washed twice with a buffer containing 5 mM HEPES-KOH, pH 7.5, 10 mM EDTA, and a cocktail of PIs. The thylakoid pellet was resuspended in 200 µl of the same buffer (Takahashi et al., 2006)

### Circular Dichroism and Melting Curves

Circular dichroism measurements were done using a spectropolarimeter (JASCO J-815) with 1 mm optical pass cuvette (Hellma). The purified recombinant proteins (100 µl) were at a concentration ≥ 100 µg/ml in Tris buffer, pH 7.5, and loaded into the clean cuvette. Single accumulation spectra ranging between 200-260 nm were recorded at RT. Scanning speed was set to 5 nm/min, with 6 sec response time and 1 nm bandwidth. Buffer blank (20 mM Tris, 10 Mm NaCl, pH 7.5) without protein served as a control for the experiment. Spectra were baseline corrected by subtracting a blank spectrum.

Melting curves were monitored using the same conditions and buffer. The single wavelength melting curve was generated at a constant wavelength of 222 nm with a temperature range (20-80°C). The CD Tool software was used to produce principal component analyses (PCA) for each sample. The two main components in the PCA analyses corresponded to spectra of folded and unfolded structures, and their magnitudes were plotted as a function of temperature, providing an overall indication of the thermal stability of the protein (CD, biopolymer).

### The PAR-PCMB Zn-Binding Assay

Zn binding by ZnJ6 was determined using the PAR-PCMB assay, as previously described (Hunt et al, 1985), except that the thiol bound zinc was released with para-chloromercuribenzoic acid (PCMB). Zinc release was measured by its interaction with 4-(2-Pyridylazo) resorcinol (PAR) at 500 nm and compared to a ZnCl_2_ standard curve. Metal-free buffers were used throughout the assay, following treatment with Chelex 100 resin (5 gr in 40 mM KH_2_PO_4_, pH 7.5), for 1 h at 37°C. ZnJ6 (3 μM) was mixed with 0.1 mM PAR in 40 mM KH_2_PO_4_ buffer to measure any free or loosely bound zinc in the solution. Addition of 30 μM PCMB to the protein solution (1 ml) caused immediate zinc release and allowed the determination and calculation of the total amount of bound Zinc per ZnJ6 molecule. PAR in buffer KH_2_PO_4_ was used as blank.

### Citrate Synthase assay

The ability to prevent aggregation of heat-sensitive proteins was tested using the Citrate Synthase assay, which monitors the holding activity of potential chaperones. ZnJ6 was added in increasing molar ratios (CS:ZnJ6, 1:0.1, 1:1, 1:2, 1:5, 1:10) to Citrate Synthase (CS) in 50 mM Tris pH 8.0 and 2 mM EDTA. CS, 12 μM is denatured by exposure to thermal stress (42°C) in a 96 well plate, containing 200 µl reaction volume in each well. The activity of ZnJ6 was measured by monitoring OD_360_ for an hour in a plate reader (BioTek Instruments, Winoosky).

### Insulin (β-chain) aggregation assay

The thiol-dependent activity of ZnJ6 was examined using the insulin turbidity assay (Arne, 1979). ZnJ6 and its cys-mutant were added to an Insulin in increasing molar ratios (ZnJ6: Insulin, 0.2:1, 0.5:1, 1:1) solution of 32 μM bovine insulin (diluted from a stock of 1.7 mM) in a freshly prepared buffer containing 0.1 M potassium phosphate (pH 7.0) and 2 mM EDTA (100 µL). The reaction was initiated by the addition of freshly prepared DTT to final concentration of 1 mM at 25°C. A reaction mix containing insulin alone served as control. Precipitation of the insulin β-chain was measured at 650 nm during 2 h, in a 96 well plate.

### Ellman’s test for determination of protein-bound sulfhydryl (PB-SH) groups

This test is used to calculate the sulfhydryl (-SH) group bound to a protein in the reaction mix. ZnJ6 was added to bovine insulin in the equimolar ratio; ZnJ6 alone served as control. One mM DTT was added to the solution to reduce the Insulin, as described above. The amount of PB-SH in the reaction mix was calculated before and after incubation of two hours. In order to quantify the PB-SH, non-protein bound -SH was subtracted from total-SH to quantify the -SH group present in the protein (Sedlak and Lindsay, 1968). Total-SH groups were quantified by adding 50 µL of reaction sample in 950 µL DTNB reagent (0.1 mM DTNB, 2.5 mM sodium acetate and 100 mM Tris, pH8). Non-protein bound -SH was measured by taking measurements after TCA precipitation. The mix was incubated for 5 min at room temperature, followed by measuring absorbance at 412 nm.

### Reduced and Denatured RNaseA Refolding Assay

Reduced and denatured RNaseA (rdRNaseA) was prepared by overnight incubation of the native enzyme (20 mg/ml) in 500 μl of 0.1 M Tris-HCl pH 8.6, containing 150 mM DTT and 6 M guanidinium hydrochloride). Excess DTT and guanidinium hydrochloride were separated from the rdRNaseA using a Sephadex G-25 buffer replacement column, equilibrated with 10 mM HCl. RNaseA aliquots (10 mg/ml stock) were stored at -80°C. Reactivation of Reduced and Denatured RNaseA was initiated by 200-fold dilution of the protein (to a final concentration of 50 μg/mL (3.8 μM) in 1mL of reactivation buffer (0.1 M Tris-HCl pH 7.0, 0.1 M NaCl and 1 mM EDTA). The refolding was performed in the absence, or presence of, at increasing molar ratios (ZnJ6: rdRNaseA, 0.2:1, 0.6:1, and 1.2:1). Aliquots (50 μL) were removed at various intervals and mixed with 50 μL of the assay mixture containing 0.1 M Tris-HCl pH 7.2, 0.1 M NaCl and 0.3 mg/ml cytidine 2′, 3′-cyclic monophosphate. RNaseA activity was measured by monitoring the hydrolysis of cytidine 2′:3′-cyclic monophosphate at 284 nm. The hydrolysis was calculated as the difference between OD_284_ at t = 0 min and t = 10 min. Refolding was presented as a percentage hydrolysis of treated samples compared to the hydrolysis of native RNaseA (Doron et al, 2018).

### Analysis of proteins that associate with ZnJ6 by pull-down experiments

Recombinant ZnJ6 fused to an SBP tag (100 μl, 10 μM) was affinity purified over streptavidin-Sepharose resin (A2S). Chloroplasts (4 ml, 0.1 μg/μl) were isolated and solubilized on ice for 5min using 1% β-DDM (n-Dodecyl β-D-maltoside, Sigma) in a buffer containing 0.7 M Sucrose, 0.1 M Tris-HCl, 0.3 M NaCl, pH 7.5 and a cocktail of PIs (Sigma). The sample was centrifuged for 40 min at 40,000 g and the soluble protein fraction was collected, diluted 1:10 in the buffer (20 mM Tris-HCl, pH 7.4, 200 mM NaCl, 1 mM EDTA), and loaded onto the streptavidin-Sepharose beads (200 μl) following their incubation with the recombinant the ZnJ6 protein. The mixture was incubated at 4°C for 2 h. The beads were then washed with 5 column volumes of the buffer (pH 7.4) to remove non-specific proteins. Finally, the bound protein with its associated complex were eluted using 2 mM biotin. SBP-tagged MBP treated similarly served as control for non-specific binding of proteins to the beads.

### Mass Spectrometry (MS)

The gel lane containing the proteins that were co-eluted from the streptavidin-Sepharose column were extracted from the gel and further reduced using 3 mM DTT (60°C for 30 min), followed by modification with 10 mM iodoacetamide in 100 mM ammonium bicarbonate for 30 min at25°C. The sample was subsequently treated with trypsin (Promega), and digested overnight at 37°C in 10 mM ammonium bicarbonate. Digested peptides were desalted, dried, resuspended in formic acid (0.1 %) and resolved by reverse phase chromatography over a 30 min linear gradient with 5% to 35% acetonitrile and 0.1 % formic acid in the water, a 15 min gradient with 35% to 95% acetonitrile and 0.1 % formic acid in water and a 15 min gradient at 95% acetonitrile and 0.1 % formic acid in water at a flow rate of 0.15 µl/min. Mass spectrometry was performed using Q-Exactive Plus mass spectrometer (Thermo) in the positive mode set to conduct a repetitively full MS scan along with high energy collision dissociation of the 10 dominant ions selected from the first MS scan. A mass tolerance of 10 ppm for precursor masses and 20 ppm for fragment ions was set. All analyses were performed in tryplicates. The MS analyses were performed in the Smoler Center in the Technion.

### Statistical analysis

Raw mass spectrometric data were analysed using the MaxQuant software, version 1.5.2.8. The data were searched against *C.reinhardtii* proteins listed in the Phytozome database. Protein identification was set at less than a 1% false discovery rate. The MaxQuant settings selected were a minimum of 1 razor/unique peptide for identification, a minimum peptide length of six amino acids, and a maximum of two mis-cleavages. For protein quantification, summed peptide intensities were used. Missing intensities from the analyses were substituted with values close to baseline only if the values were present in the corresponding analysed sample. LFQ intensities were compared among the three SBP-ZnJ6 biological repeats and the three SBP-MBP repeats on the Perseus software platform using the student t test.

The enrichment threshold (LFQ intensity of SBP-ZnJ6 subtracted from SBP-MBP control) was set to a log2-fold change ≤ -3 (8 fold enrichment as compared to control) and p-value < 0.05. The filtered proteins were categorised both manually, based on their function in the Phytozome database, and using BLAST2GO software based on the biological process (with a minimum of 2 fold enrichment) as selected criteria. The minor categories (sub-branches) with the BLAST2GO software were merged to get the four broader classes (photosynthesis, metabolic process, oxidoreductases, and transporters)

### Redox sensitivity of *Chlamydomonas* cells by their exposure to H_2_O_2_, MeV or DTT

To verify the *in vivo* function of ZnJ6, the insertional knock-down mutant with the paromomycin resistance was used. *C. reinhardtii* cells (mutant, wild type background cells) were grown to mid-log phase in HS medium with light intensity of 150 µmol/s/m^2^. The mutant was confirmed by PCR and western analysis using anti-ZnJ6 antibodies. For all the treatments, ∼1× 10^7^ cells ml^−1^ (1 ml) were exposed to different concentrations of H_2_O_2_ (2, 5, 10, and 20 mM), MeV (2, 5, 10, and 20 µM) and DTT (2, 5, 10 and 20 mM). After exposure, the cells were washed twice and resuspended in HS media with required dilution. 10^3^ cells (3 µl) were seeded over HS plates and allowed to grow at 23°C. Pictures were taken on day 5 of the growth and analysed using the MultiGauge software.

### Exposure of *Chlamydomonas* cells to heat stress

*C. reinhardtii* cells (mutant, wild type background cells) were grown as explained earliers (in Chlamydomonas strains and Growth Condition section). 5 µl (serially diluted from 10^5^ to 10^2^ cells/ 5 µl) aliquots were spotted on the HS plates and incubated at 23 °C (control) or exposed to heat stress by incubating the plates at 37 °C for 1, 2, and 4 h. Plates were then returned to 23°C and allowed to grow for an additional 5 days. A loop full (1 µl inoculation loop) of cells from 10^5^ cell lane was then taken and resuspended in 100 µl of HS media, 5 µl of cells were spotted on a new plate and allowed to further grow at 23°C for another 5 days to check the growth rate.

## Supplemental Data

**Supplemental Table S1**. Mass spectrometry analysis of pulled down proteins.

**Supplemental Table S2**. Primers for recombinant protein expression.

**Supplemental Table S3**. Primers for Cysteine to Serine (TGC > TCC) mutagenesis of ZnJ6.

**Supplemental Methods-** Cloning for recombinant protein expression.

**Supplemental Figure S1**. ZnJ6 antibody generation.

**Supplemental Figure S2**. Prediction of the transmembrane domain of ZnJ6

**Supplemental Figure S3**. Purified recombinant protein run on SDS-PAGE

**Supplemental Figure S4**. Percoll two-step gradient for chloroplast isolation and schematic representation of pull-down experiment along with anti-SBP western analysis of different pull-down fractions.

**Supplemental Figure S5**. Characterization and verification of CLiP mutant LMJ.RY0402.048147.

## ACKNOWLEDGEMENTS

We are grateful to Prof. Zach Adam for antibodies against OEE33 and Moshe Sagi for antibodies against psbA. The GB1 tag for protein stability was obtained from Prof. Gerhard Wagner. We thank Prof. Dudy Bar-Zvi, Prof. Ofer Yifrach and Prof. Michele Zaccai for valuable comments. We also thank Dr. Lior Doron, Dr. Naama Segal, Dr. Irit Dahan, Nofar Baron, for helpful discussions, and Dr. Matan Drory for helping with BLAST2GO enrichment analysis.

